# Comprehensive analysis of microbial content in whole-genome sequencing samples from The Cancer Genome Atlas project

**DOI:** 10.1101/2024.05.24.595788

**Authors:** Yuchen Ge, Jennifer Lu, Daniela Puiu, Mahler Revsine, Steven L. Salzberg

## Abstract

In recent years, a growing number of publications have reported the presence of microbial species in human tumors and of mixtures of microbes that appear to be highly specific to different cancer types. Our recent re-analysis of data from three cancer types revealed that technical errors have caused erroneous reports of numerous microbial species found in sequencing data from The Cancer Genome Atlas (TCGA) project. Here we have expanded our analysis to cover all 5,734 whole-genome sequencing (WGS) data sets currently available from TCGA, covering 25 distinct types of cancer. We analyzed the microbial content using updated computational methods and databases, and compared our results to those from two major recent studies that focused on bacteria, viruses, and fungi in cancer. Our results expand upon and reinforce our recent findings, which showed that the presence of microbes is far smaller than had been previously reported, and that many species identified in TCGA data are either not present at all, or are known contaminants rather than microbes residing within tumors. As part of this expanded analysis, and to help others avoid being misled by flawed data, we have released a dataset that contains detailed read counts for bacteria, viruses, archaea, and fungi detected in all 5,734 TCGA samples, which can serve as a public reference for future investigations.

**One-sentence summary:** Analysis of microbial content in 5,734 whole-genome sequencing samples from TCGA yields a comprehensive new resource for investigating the role of microbes in cancer.

## Introduction

A number of recent studies have used the vast sequencing resource created by The Cancer Genome Atlas (TCGA) project to explore the potential role of microbial species in cancer. Although most of the TCGA data was collected with the goal of studying human genetic variation or gene expression, microbes present in the tumors–including viruses, bacteria, and fungi–might also be captured as an incidental side effect of sequencing experiments. Identifying microbes in a human tumor sample, in which the vast majority of the biomass is expected to be human in origin, requires great care in order not to be misled by contaminants, sequencing vectors, or other artifacts that might also be present in the data. In this study, our objective was to conduct an exhaustive and meticulous survey of microbial communities across thousands of whole-genome sequencing (WGS) samples from the TCGA project, with the goal of identifying any microbes within these samples. By making our results publicly available, we hope to spur additional research that may amplify or alternatively refute recent findings of microbiomes in a wide variety of tumor types.

We also compare our findings to two recent studies that used much of the same TCGA data and described findings that were, in some instances, substantially affected by contamination. Those studies and others that have relied on their data have implicated the microbiome in various aspects of cancer, from modulating the tumor microenvironment to influencing treatment responses. In the first study, Poore *et al. (1)* analyzed 17,625 samples from TCGA, and reported that they were able to use machine learning algorithms to construct highly discriminative microbial signatures in 32 of 33 types of cancer. Their classifiers, which used combinations of bacteria, archaea, and viruses, were remarkably accurate, obtaining 95-100% accuracy at discriminating each of the 32 cancer types against all others. They reported additional models with similar accuracy at distinguishing tumors from matched normal samples in 15 cancer types. (Note: due in part to our findings *(2)*, the journal *Nature* retracted the Poore *et al*. study in July of 2024 *(3)*.) In the second study, Narunsky-Haziza *et al. (4)* analyzed 17,401 samples from the TCGA project and other sources with the goal of identifying associations between cancer and fungi, which the first study had not considered. They reported distinctive fungal signatures of cancer in most of the 35 cancer types they considered *(4)*. Our analysis here looks at many of the same TCGA samples in an effort to replicate some of these findings.

One source of over-counts when analyzing human samples for microbial content is data contamination in public genome databases. As described previously, the inadvertent inclusion of human DNA within microbial genomes has affected thousands of genomes *(5)*. When creating a microbial sequence database, it is crucial to be aware of this issue and to take rigorous steps to remove these computational contaminants, which otherwise may substantially skew the results of metagenomic analyses. Such contamination events can be especially problematic when working with low biomass samples, where the microbial content is expected to be a very small proportion of the total sample. This is precisely the scenario encountered when “mining” human DNA sequencing projects such as TCGA for microbial content.

We should emphasize further that even a tiny amount of contamination in a genome’s database sequence can lead to enormous over-counts of bacterial species, for the following reason. The most common source of human contamination in bacterial genome databases is high-copy human repeats such as Alus, LINEs, and SINEs, as described earlier *(5)*. Any WGS sample from human tissue is likely to contain large numbers of reads from these widespread repeats. Thus if one scans a human DNA sample against a bacterial genome that is contaminated with even one of these human repeats, many human reads will appear to match the bacterium. For a typical TCGA sample, it would not be surprising to find tens or even hundreds of thousands of reads incorrectly matching a bacterial genome in this circumstance *(2)*.

Vector contamination, in which reads deriving from vectors such as manufacturer-specific sequencing primers make their way into a genome assembly, further compounds the challenges associated with metagenomic analyses. Sequences originating from vectors have inadvertently found their way into genome databases, where they might be labeled as bacteria, fungi, plants, or animals. As we describe below, some fungal genome sequences in public databases are contaminated with vector or adaptor sequences, which can lead to large numbers of false positive matches to a sample that was sequenced using the same vectors.

## Results

We analyzed the microbial content of 5,734 WGS samples from TCGA, which comprised all of the available WGS samples as of late 2023. Despite the fact that TCGA also includes large numbers of RNA sequencing (RNA-seq) experiments, we excluded them because they used poly-A selection to capture messenger RNA. Bacterial transcripts do not have long poly-A tails

*(6)* and will not be captured, except very rarely, with poly-A selection protocols. Thus any bacterial sequences found in a human RNA-seq experiment are almost certain to be contaminants, and the inclusion of human RNA-seq data in a search for bacterial signatures, as has been done occasionally in previous studies *(1)*, simply does not make sense.

We removed human sequences from the TCGA data by mapping the reads against both the GRCh38 reference genome and the CHM13 human genome (see Methods). As shown in **Table 1**, after removal of human sequences, the number of reads remaining in most samples was relatively small, averaging 2.6 million reads per sample (0.48% of the total). Across all samples, a total of 15 billion reads remained after two-pass filtering. Of these, we identified 2.44 billion as vector contaminants during classification with Kraken.

**Table 1.**
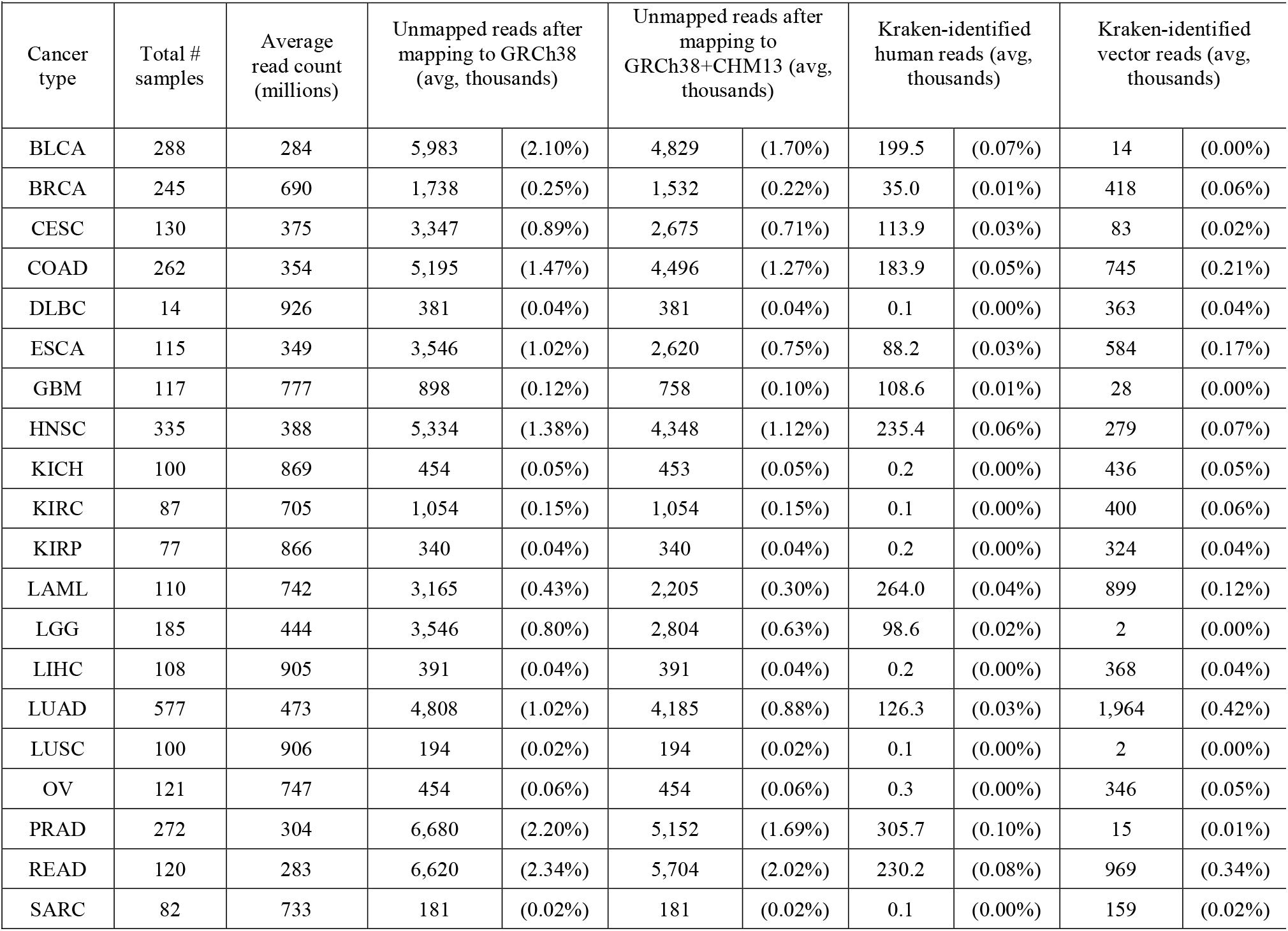
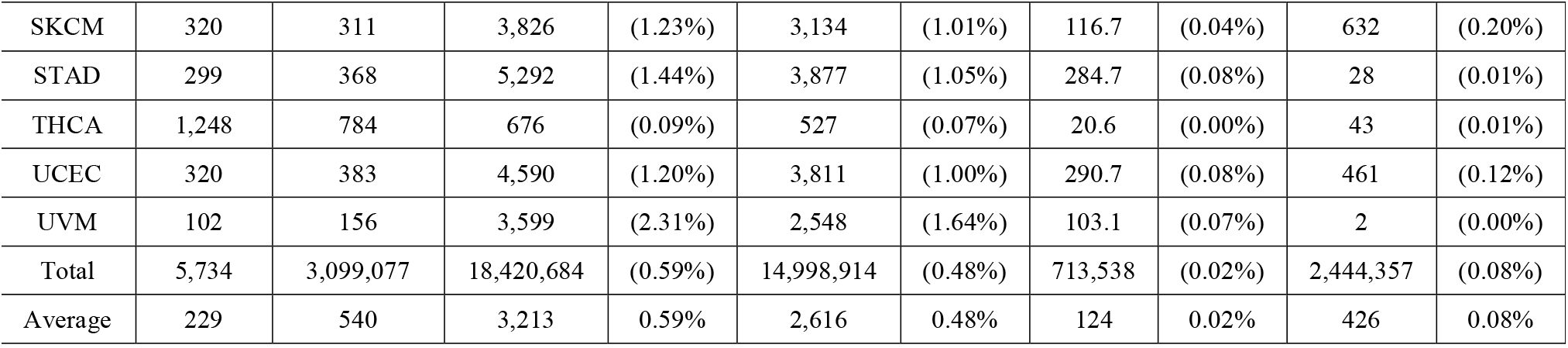
Overview of the 5,734 WGS samples analyzed in this study. Column 4 shows how many reads remained per sample after removing reads that mapped to the GRCh38 human genome. Column 6 shows the number of reads remaining after a second pass further removed reads matching the CHM13 genome. Columns 8 and 10 show the average numbers of reads from column 6 that were identified by Kraken as either human or vector.

We used KrakenUniq *(7)* to classify the filtered reads against a microbial database containing 50,651 genomes representing 30,355 species including bacteria, archaea, viruses, and fungi (see Methods). Across all samples, we identified 11,332 non-eukaryotic microbial species that occurred at least once. **Table S1** reports the number of sequencing reads detected for every species in each of the samples. **Table S2** includes normalized counts for the data in **Table S1**, where read counts are divided by the total number of reads sequenced for each sample, in millions. **Table S3** further normalizes the counts by dividing by the genome size of each species, in kilobases. The vast majority of the entries in these tables are zero: out of ∼65 million entries, only 1,881,550 (2.9%) have non-zero values, and only 214,325 (0.3%) have raw read counts of 10 or greater. We then classified reads that were not identified in the previous step against a comprehensive fungal genome database containing 557 species (see Methods). **Table S4** reports the raw read counts for all 5,734 samples for each of these fungal species, with normalized counts reported in **Tables S5-S6**.

Note that a small number of viruses had already been removed from the samples by the TCGA project, which used bwa *(8)* to align all reads to multiple subtypes of human papillomavirus, hepatitis B and C, and cytomegalovirus. Because these reads were identified by bwa and not KrakenUniq, we provide read counts for those species separately in **Table S7**.

Worth noting here is that even after alignment against two human genomes, an average of ∼124,000 reads per sample were still classified as human by the Kraken program (**Table 1**). This highlights the necessity of including the human genome in any metagenomics database, even if the data are pre-filtered to remove human reads. **Figure 1** shows the breakdown of bacterial, viral, and fungal read counts across the 25 cancer types, whose abbreviations are described in **Table S8**.

**Figure 1.**
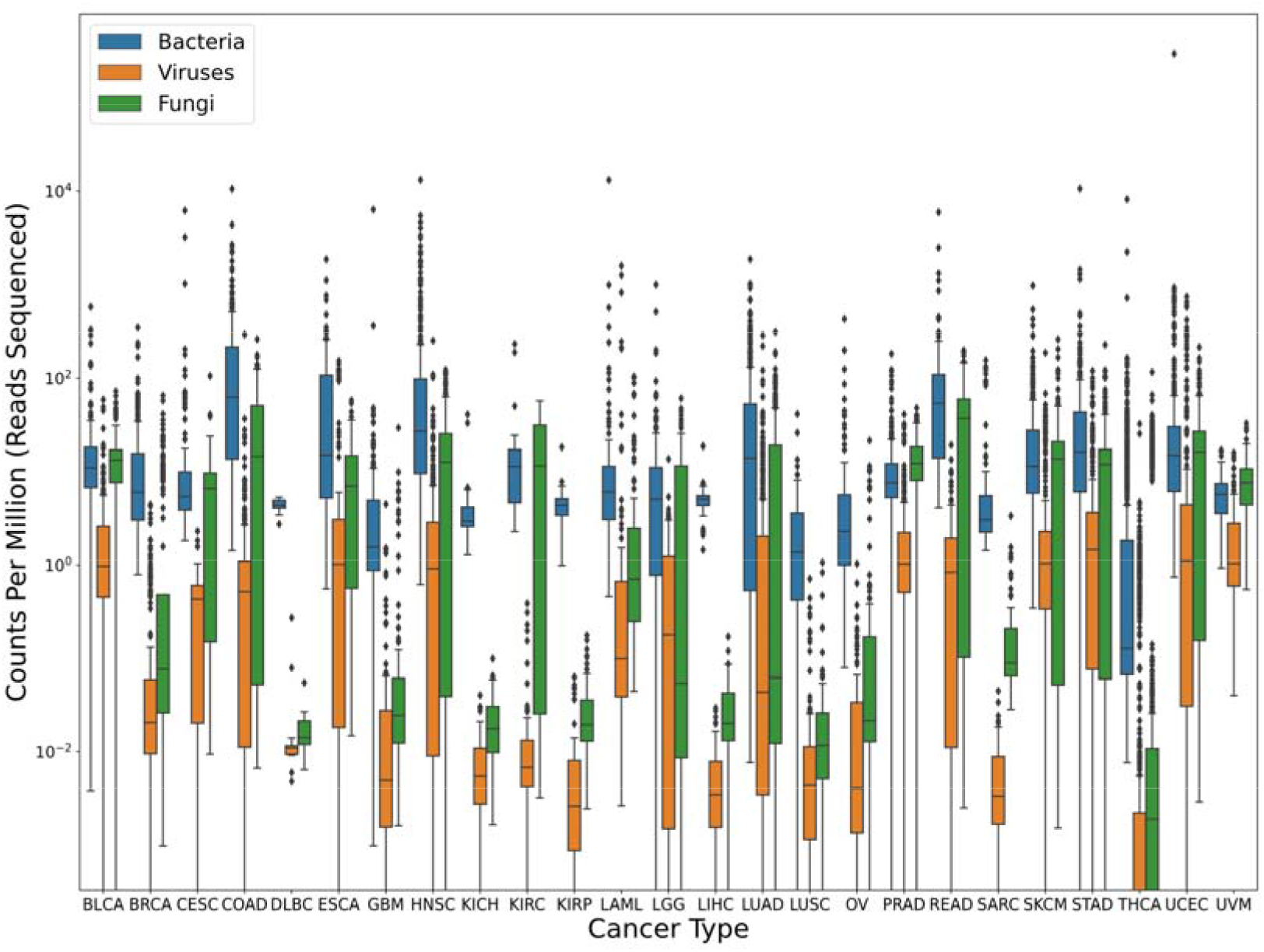
Microbial content found in TCGA data from 25 cancer types. Box plots show the inter-quartile range of read counts for the samples from each cancer type, with a horizontal line indicating the mean value. On average, bacteria (blue) had much higher read counts than fungi (green), or viruses (orange). Read counts were normalized by the number of reads sequenced in each sample, in millions; i.e., a value of 10 indicates 10 reads for every million reads in the sequencing run. Note that no adjustments were made for genome size here, which explains the much smaller read counts for viruses. For clarity, the y-axis uses a log scale.

Interestingly, a number of samples showed unusually high read counts for a single pathogen, which might in some cases indicate an infection. For example, although only a handful of samples contained more than 100 reads from *Helicobacter pylori*, one sample contained >175,000 reads. That sample was from stomach cancer (STAD) data but the source was normal tissue rather than a tumor. For *Fusobacterium nucleatum*, which has been associated with colon cancer, only 123 samples had >100 reads, but three samples had >100,000 reads with one having 372,665 reads, found in a tumor sample from head and neck cancer (HNSC).

A previous study of TCGA data reported that the cervical cancer (CESC) dataset had a high proportion of samples positive for human papillomavirus, particularly HPV-16 and HPV-18 *(9)*. Although the earlier study relied on PCR for detection, which should be more sensitive than whole-genome sequencing, our analysis of the WGS data confirmed their findings. Among the 66 non-tumor CESC samples, nearly all had very low counts for all subtypes of HPV, with only one sample containing >1000 reads for HPV-16. In contrast, among the 64 CESC tumor samples, 31 samples had >1000 reads from HPV-16, 8 other samples had >1000 reads from HPV-18, and 11 more had >1000 reads from other HPV strains (**Table S7**). Also worth noting is that 33 of 171 tumor samples in head and neck cancer (HNSC) had >1000 reads from HPV-16, while all of the normal samples from HNSC had fewer than 200 HPV-16 reads.

### Abundant bacteria, viruses, and fungi may be contaminants

The most abundant species across all 5,734 samples, in our analysis, were *Delftia acidovorans, Rothia mucilaginsa, Human mastadenovirus C, Pseudomonas sp. J380, Escherichia coli, Bacteroides fragilis, Prevotella intermedia*, and other Delftia and Rothia species. Each of these i known to be present on human skin, in the oral cavity, or in the gut, and each was found in large numbers of both tumor and normal samples, ranging from 378 samples containing *Human mastadenovirus C* to 1,968 samples with *D. acidovorans* and 4,440 samples with *E. coli*. The mundane and perhaps most likely explanation for the presence of these species, especially for tissue samples from internal organs not known to harbor a microbiome, is that most of these findings represent contamination, with possible sources including the sequencing facilities and the multiple people handling each of the samples. For example, the most frequently observed fungal species, appearing in 2,312 samples, was *Saccharomyces cerevisiae*, a widely-used model organism that is not a human pathogen and that frequently appears as a cross-contaminant in sequencing centers. Another example is the bacterium *Cutibacterium acnes*, a common human skin bacterium that was abundant in most of the cancer types. Another abundant species was *Rosellinia necatrix partitivirus 8*, a virus that infects plant fungi and has no known associations with human disease *(10, 11)*.

### Associations between microbes and cancer types

To determine if any of the microbial species detected in our analysis might be associated with cancer, we examined the most abundant microbes in each of the cancer types, using the normalized values in **Table S3**, and also looked specifically for bacteria and viruses that have known associations with cancer. The top species for each cancer type are shown in **Supplementary Figure S1**.

We noted that after normalization, a few known cancer-associated microbes appeared relatively abundant; e.g. as mentioned above, HPV was identified in most of the cervical cancer samples. We also found that *Bacteroides fragilis* was the top species for rectal adenocarcinoma (READ) and the fourth-most abundant microbe in colon adenocarcinoma (COAD), and *H. pylori* was the 17^th^-most common microbe in stomach adenocarcinoma (STAD).

A small number of abundant species might represent acute infections from the original patient samples. *Metamycoplasma salivarium*, an oral pathogen that may be pathogenic in immunocompromised individuals, was the top species in esophageal carcinoma *(12)*, and this finding might merit follow-up investigation. Although we found no evidence to support claims of a microbiome–i.e., a community of microbes–in any of the cancer types, the datasets provided here contain hundreds of species that we did not investigate, and that might be worthy of further study.

### Comparison of bacterial and viral read counts to previous reports

In a recent re-analysis *(2)* of TCGA sequence data from three cancer types–bladder cancer, breast cancer, and head and neck cancer–we reported that an earlier study (now retracted *(3)*) by Poore *et al*. of the same data *(1)* had described read counts that were far too high. In particular, we found that >95% of the read counts in the earlier study were inflated by at least a factor of ten *(2)*. Using the more-comprehensive data here, we can now confirm the previous findings and extend them to all 25 cancer types for which WGS data is available.

The Poore *et al*. study *(1)* reported read counts for 1,993 microbial genera, including bacteria, archaea, and viruses. Out of the 5,734 WGS samples analyzed in this study, we identified 4,550 samples that matched those from the Poore *et al*. study; the remaining samples were added to TCGA more recently and thus were excluded from this comparison. Our analysis found reads from 2,852 genera across the 4,550 samples, which included 1,285 found in the previous study. The union of the two sets yielded 3,552 genera found in one or both studies (**Figure 2**). (Note that we excluded the genera for HPV, HBV, HCV and CMV from this figure because those viruses, shown in **Table S7**, were pre-filtered by TCGA).

**Figure 2.**
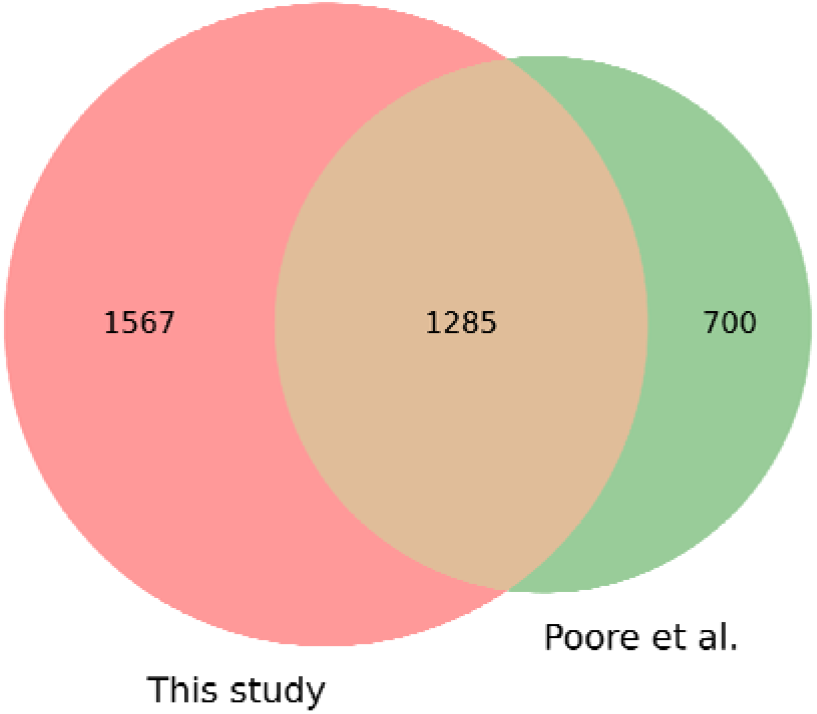
Comparison of bacterial, viral, and archaeal genera found in 4,550 TCGA samples versus those found in an earlier study. We found a total of 2,852 genera while a previous study ^1^ found 1,993, with 1,285 found in both studies.

Supplementary **Tables S9** and **S10** contain read counts for all 4,550 samples and for the 3,552 genera found in either study, with **Table S9** containing the results for this study and **Table S10** showing the corresponding counts from Poore *et al*. We compared the findings by computing the ratio of counts (**S10/S9**), replacing any zero values with 1 to avoid division by zero. We then examined the ratios of read counts for all cells with a count ≥10 in at least one study, which comprised 2,189,351 entries.

This analysis showed that the microbial read counts reported by Poore *et al*. were vastly higher than the counts found in here. The median ratio was 56; i.e., half of the read counts in the earlier study were at least 56 times too high, and 90% of the values were at least 11 times too high. As shown in **Figure 3**, the top 5% (109,472) of the Poore *et al*. read counts are more than 9,282-fold too high. The primary reason for these extreme over-estimates, as explained previously *(2)*, was the use of a database containing thousands o incomplete (“draft”) bacterial genomes, which themselves were contaminated with human sequences. As a result, millions of human reads in the TCGA data were mistakenly identified by Poore *et al*. as bacterial reads.

**Figure 3.**
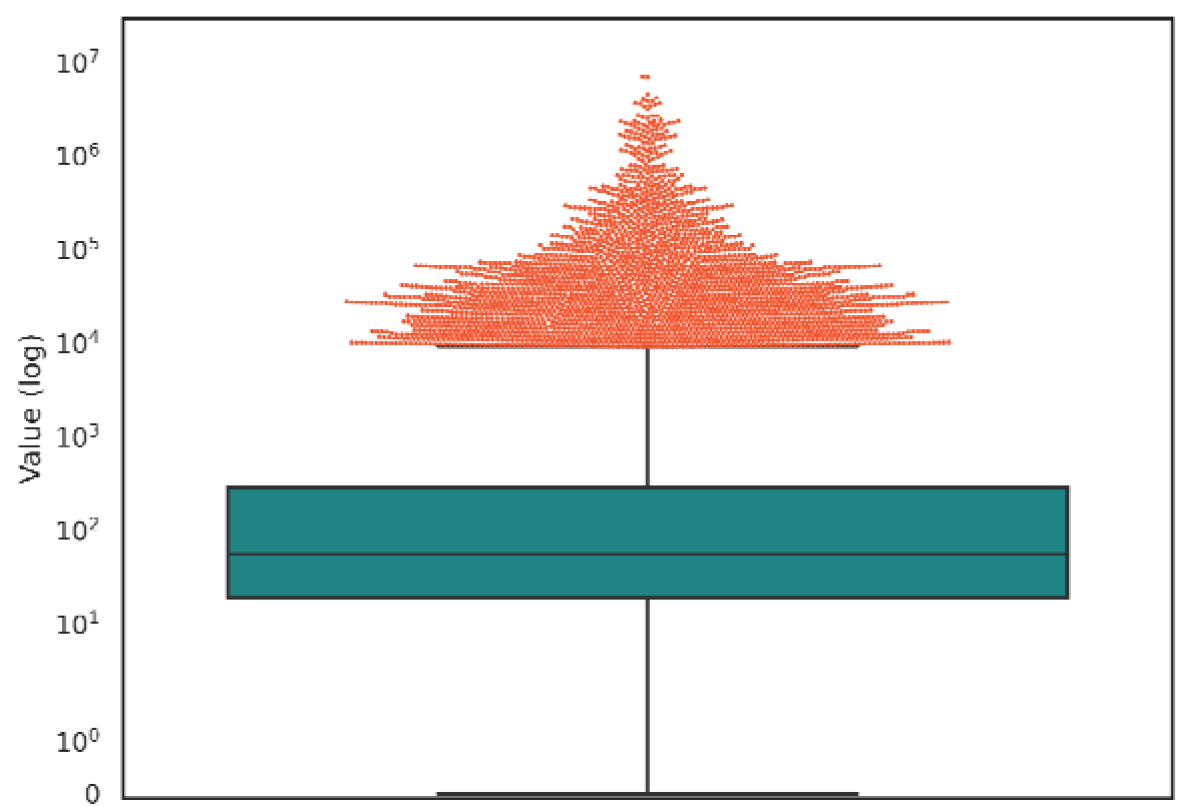
Distribution of ratios between read counts found by Poore et al. (Poore et al. 2020) and this study, for all samples in which either study found at least 10 microbial reads. The boxplot shows the median (56) and interquartile range. The top 5% of points, comprising 109,472 values, are shown as a cloud above the value 9,282.

To provide another comparison, we identified the top 10 most-abundant microbial genera across all 4,550 WGS samples reported in the earlier study and compared them to our read counts for the same genera. As shown in **Figure 4**, the top three genera in Poore *et al*. were *Streptococcus, Mycobacterium, and Staphylococcus*, with average read counts per sample of 1,780,000, 1,400,000, and 922,000, respectively. In our re-analysis of the same samples, we found average read counts of 1129, 31, and 39 reads in those genera, values that range from 1,500 to 45,000 times smaller.

**Figure 4.**
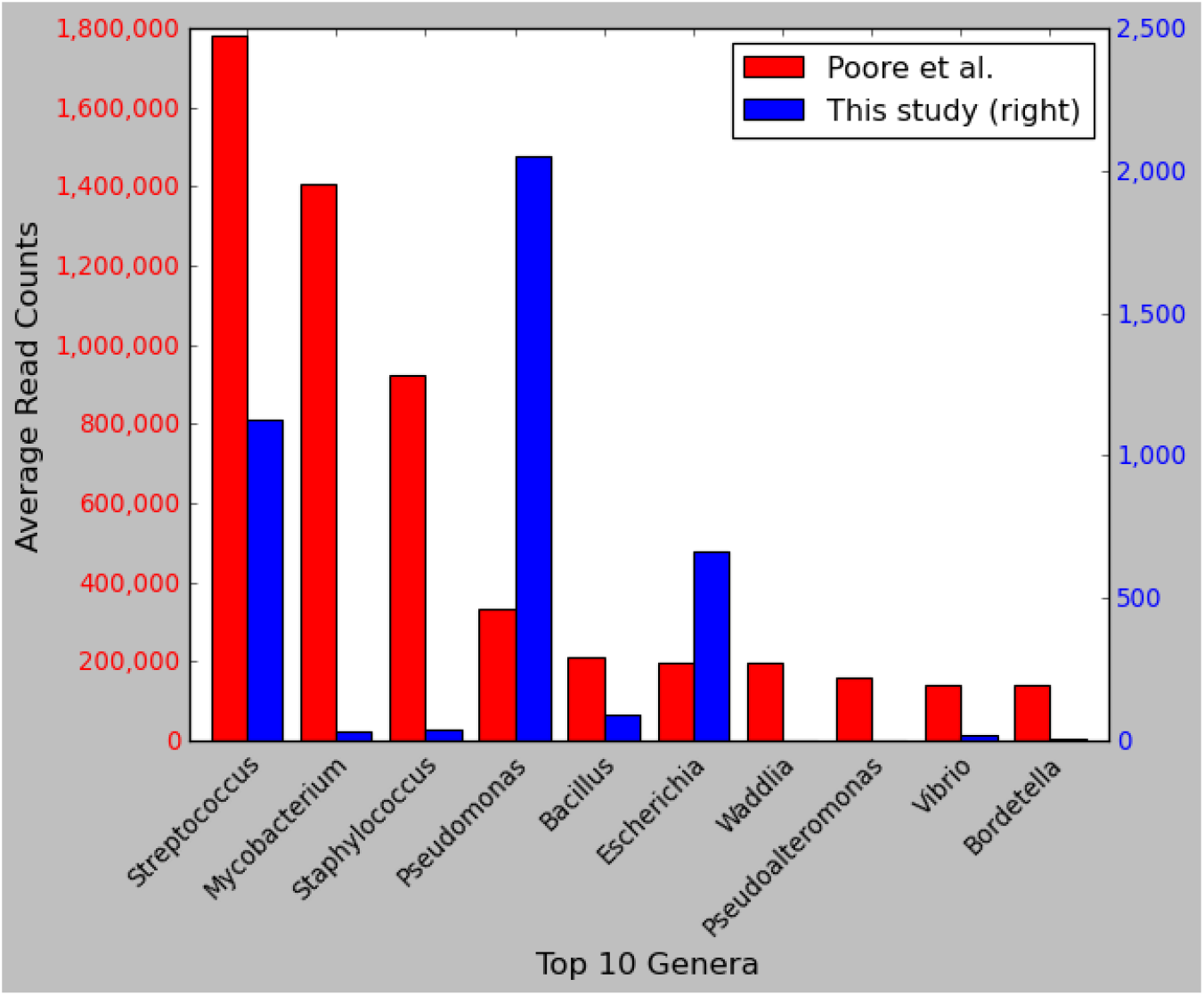
Comparison of read counts for the top 10 microbial genera from a previous study. Shown are the top ten most-abundant genera ranked by the average number of reads reported in Poore *et al. (1)* across 4,550 WGS samples from 25 cancer types. The left axis shows read counts as reported by Poore *et al*., and the right axis shows read counts as computed in our re-analysis of the same samples and the same species. Note that the y-axis scales differ by a factor of ∼700. The x-axis shows genus names.

Only 2% of the values in our study were higher than those in the previous study. Among the 1,567 genera found exclusively in our study, the average read count per sample was just 0.81, suggesting that most are either false positives or low-level contaminants. The most abundant species found exclusively in our analysis was *Schaalia turicensis* (previously called *Actinomyces turicensis (13)*), a bacterium that is commonly found in the oral and gut microbiome.

### Comparison of fungal content to previous reports

In another recent study using the TCGA data, Narunsky-Haziza *et al. (4)* reported a strong association between mixtures of fungal species, which the authors called a “mycobiome,” and multiple cancer types. Out of the 5,734 WGS samples analyzed here, we identified 4,271 that were identical to those used in the Narunsky-Haziza *et al*. study. We re-analyzed these samples for fungal content using a separate fungal genome database that contained 557 species, including all 224 of the species included in the Narunsky-Haziza study (see Methods). The full set of read counts for the 4,271 samples used in both studies can be found in Supplementary **Table S11** and **S12**, where the counts in **Table S12** were taken from Narunsky-Haziza *et al. (4)* Note that because we used a superset of the fungal species, we identified reads from many species not reported in the previous study; however, these were observed at very low levels, with an average read count of 3.8 reads per sample in species unique to our analysis.

Although our average read counts were in rough agreement, we found highly divergent results for a small number of species that were estimated by Narunsky-Haziza *et al*. to be highly abundant. These are illustrated in **Figure 5**, which compares the maximum read counts from any sample for the top 10 most-abundant fungal species from Narunsky-Haziza *et al*. These include samples containing 2,013,180 reads from *Ramularia collo-cygni*, 656,503 reads from *Trichosporon asahii*, 101,344 reads from *Candida albicans*, and 54,641 reads from *Malassezia globosa*. In contrast, our counts for the same samples were 332, 4622, 266, and 4, values that range from 142-fold to 13,660-fold smaller.

**Figure 5.**
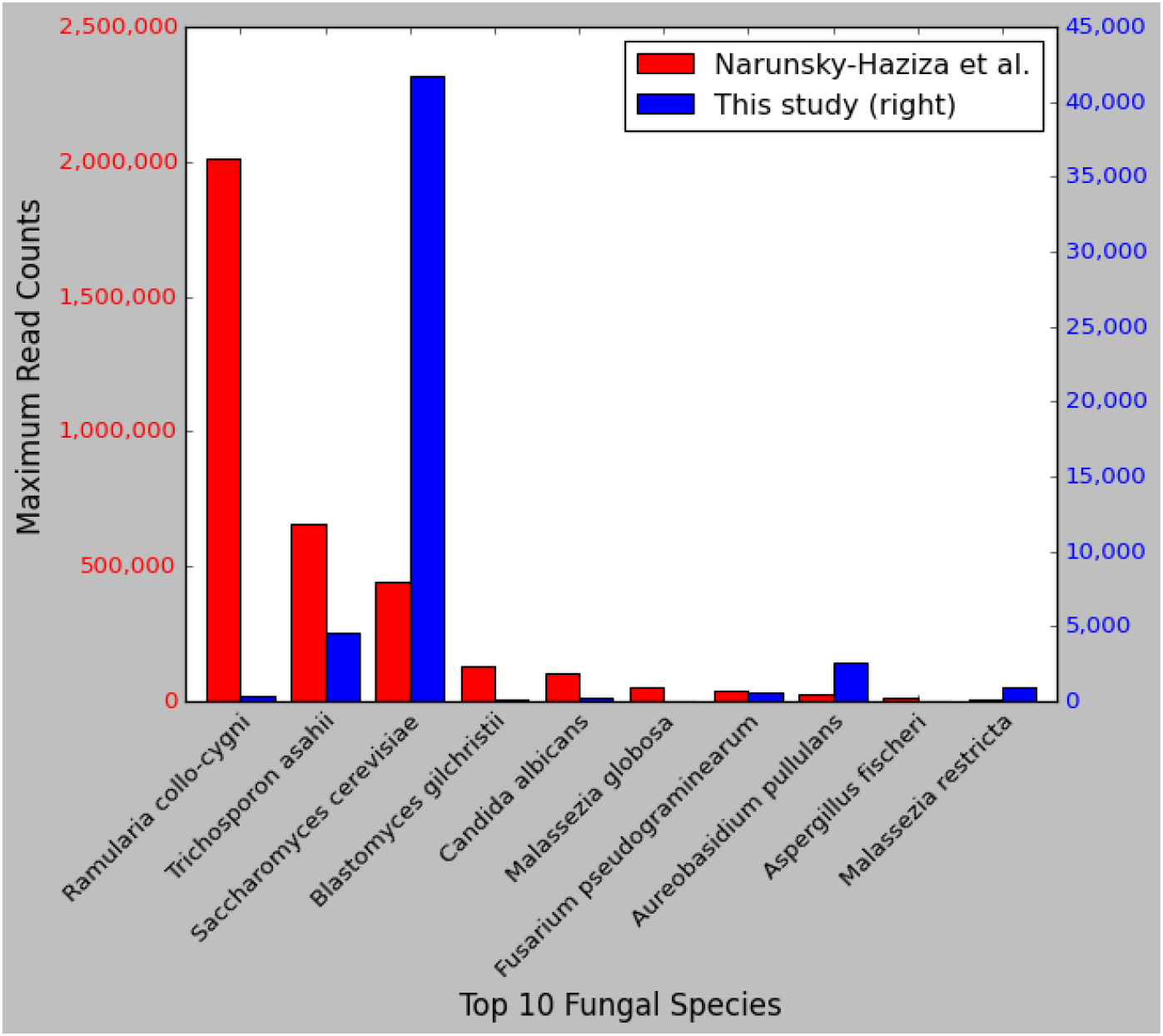
Comparison of read counts in fungal species. Shown are the top ten most-abundant fungal species ranked by the maximum number of reads reported in Narunsky-Haziza *et al. (4)*, across 4,271 WGS samples from 25 cancer types. The left axis shows read counts as reported by Narunsky-Haziza *et al*., and the right axis shows read counts as computed in our re-analysis for the same samples and the same species. Note that the y-axis scales differ by a factor of 50. Note also that Narunsky-Haziza *et al*. reported a maximum of ∼441,000 reads from one sample matching *S. cerevisiae*, a finding that we were unable to replicate. In our analysis of the same sample (see Methods), we found ∼40,000 reads matching *S. cerevisiae*, as shown in the figure. These matches predominantly fell within a region of the yeast genome that contains the 18S rRNA gene and that has significant similarity to human, suggesting that these were human reads that slipped past the filtering step.

Notably, the species with these large read counts were considered particularly important by Narunsky-Haziza *et al*., and were used by them to define three fungi-driven “mycotypes,” labeled F1 (*Malassezia, Ramularia*, and *Trichosporon*), F2 (*Candida, Aspergillus*), and F3 (multiple genera including *Yarrowia*), which they reported were associated with distinct immune responses and overall survival *(4)*. Below we explain how contamination in the genomes themselves led to at least some of the higher read counts. We hypothesize that if the raw counts are corrected, these mycotypes and their association with cancer may be substantially reduced.

### Contamination in the *Malassezia globosa* genome

The highest read count for *M. globosa* as reported by Narunsky-Haziza *et al*. was 54,641 reads from sample h2540, a blood-derived normal sample from the head and neck cancer (HNSC) dataset, in which our re-analysis found only 4 *M. globosa* reads. We subsequently aligned all reads (without filtering) from sample h2540 against the *M. globosa* reference genome (see Methods) and found a very large number of matches, nearly all aligning to just two locations: an 897-bp contig and a 557-bp contig (**Table 2**). We then used BLAST *(14, 15)* to confirm that both contigs were human sequences that were mis-labeled as *M. globosa*. Note that the *M. globosa* genome assembly was revised in December 2023 (GenBank accession GCF_000181695.2), and the contigs shown in **Table 2** were removed by NCBI because they were determined to be contaminated.

**Table 2.** Contaminants found in fungal genomes in GenBank. Columns 1-3 show the species name, GenBank accession number of the genome, and accession number of the scaffold in which the contaminant was found. Columns 4-6 show the starting location of the contaminant within the scaffold, its length, and the correct source of the DNA sequence.

| Organism | Accession | Scaffold Accession | Start | Length | Contamination Source |
| --- | --- | --- | --- | --- | --- |
| *M. globosa* | GCF_000181695.1 | NW_001849834.1 | 1 | 897 | Human |
| *M. globosa* | GCF_000181695.1 | NW_001849877.1 | 1 | 557 | Human |
| *T. asahii* | GCF_000293215.1 | NW_014040855.1 | 924,646 | 58 | Vector: NGB00360.1, Illumina PCR Primer |
| *T. asahii* | GCF_000293215.1 | NW_014040868.1 | 203,990 | 64 | Vector: NGB00852.1, NEBNext Index 6 Primer |
| *R. collo-cygni* | GCF_900074925.1 | NW_019716264.1 | 394,971 | 58 | Vector: NGB00360.1, Illumina PCR Primer |
| *R. collo-cygni* | GCF_900074925.1 | NW_019716256.1 | 1,478,660 | 58 | Vector: NGB00360.1, Illumina PCR Primer |

### Contamination in the *Trichosporon asahii* genome

*Trichosporon asahii* also had an unusually high read count in the Narunsky-Haziza *et al*. study, which reported a maximum count of 656,503 reads in sample h1948 (a solid tissue normal sample from TCGA-LUAD). Upon aligning all reads (unfiltered) from that sample to the *Trichosporon* genome, we found an even higher number of matches, over 54 million; however, 99.99% of the matches hit the same 80-bp interval in the genome. We investigated and found that 58bp from this 80bp sequence was identical to an Illumina sequencing vector (accession NGB00360.1); i.e., it is a contaminant in the genome assembly. The vector contaminant occurs in the middle of a large, 2.7 Mbp scaffold (NCBI accession NW_014040855.1, see **Table 2**).

Also worth noting here is that sample h1948 was a failed sequencing run, in which 95.4% of the read pairs (160 million out of a total of 179 million) were vector.

We found similar results for other samples; e.g., Narunsky-Haziza *et al*. reported 302,882 *Trichosoporon* reads in sample h1325, a lung cancer tumor sample. When we aligned the entire set of reads (unfiltered) from this sample to *T. asahii*, we found 12,235,713 matching reads, and all except for 18 reads aligned to the same 58-bp vector contaminant mentioned above. In a different sample, we found 1980 reads matching a 64-bp interval in the *T. asahii* genome that turns out to be another vector (accession NGB00852.1), also shown in **Table 2**. Thus for these and other samples, the large numbers of apparent matches to *Trichosporon* appear to have been the result of vector contaminants in the fungal genome sequence.

### Contamination in the *Ramularia collo-cygni* genome

Sample h1948, a normal tissue sample from the lung cancer (LUAD) dataset, was reported by Narunsky-Haziza *et al*. to contain 2,013,180 reads from *R. collo-cygni*, whereas we found only 322 reads. We re-aligned the filtered reads to the *R. collo-cygni* genome using the same Bowtie2 parameters as used by Narunsky-Haziza *et al*., and found ∼34M matching reads, all matching just two locations: a 77-bp sequence on RCC_scaffold10 (NCBI accession NW_019716264.1) and a 77-bp sequence on RCC_scaffold02 (NCBI accession NW_019716256.1, see **Table 2**). Both of these sequences contain a 58-bp subsequence identical to an Illumina PCR primer, one of the two that we found in *Trichosporon*. Thus these apparent matches to *Ramularia* appear to have been the result of vector contamination.

### Analysis of high read counts for *Candida albicans*

The highest read count reported by Narunsky-Haziza *et al*. for *C. albicans* was 101,344 reads from sample h949, a primary tumor sample from the head and neck cancer (HNSC) dataset. We attempted to replicate this finding by processing the sample using the host depletion pipeline described in the original study, which left only 93,556 reads, a number already lower than the reported count for *C. albicans*. We aligned these reads to the *C. albicans* genome using Bowtie2 *(16)*, which detected only 230 matching reads. Using the same procedures, we analyzed the sample with the second highest *C. albicans* read count, h5103, a normal tissue sample from the lung cancer (LUSC) dataset with 75,799 reported matches, and found only 544 matching reads. We were similarly unable to replicate high read counts for *C. albicans* in other samples.

To attempt to explain the far higher read counts reported by Narunsky-Haziza *et al*., we aligned the entire set of unfiltered reads (i.e., without first removing human-matching reads) from the HNSC and LUSC samples to the *C. albicans* genome, which yielded 75,392 and 133,576 matching reads, respectively, values closer to the original report. However, 91% of the 75,392 reads from the HNSC sample were originally mapped to human in the raw TCGA data, and thus should not have been included in the “non-human” read sets. We investigated further and determined that nearly all these reads matched ribosomal RNA (rRNA) genes in both human and *C. albicans*, although the matches to *C. albicans* were far better. This analysis suggests that *C. albicans* was genuinely present in the samples (whether it was in the tissues or a contaminant), but it is unclear how the pipeline in the original study yielded these high read counts.

## Discussion

Using updated computational methods and databases, we have analyzed the non-human content in a large collection of whole-genome sequencing data sets in TCGA covering 25 distinct types of cancer, and created a comprehensive dataset encompassing detailed read counts for bacteria, viruses, archaea, fungi, and other microbes in 5,734 samples from tumors and normal tissue. Our data show that most of the read counts reported in a previously-published dataset are greatly inflated, often exceeding the true counts by factors exceeding 1,000. As we explained previously *(2)*, these over-counts can be attributed in part to the inclusion of numerous draft bacterial genomes (which themselves contain contaminants) in the database used for the metagenomic analysis. Although the primary paper was recently retracted, the results from Poore *et al. (1)* have been used in at least a dozen other studies *(17–28)* that downloaded their read count data and then published findings based on that data, and another recent study *(29)* similarly based its results on the “mycobiome” data from Narunsky-Haziza *et al. (4)*. Our analyses here used a cleaner genome database, which yielded much lower read counts for the microbial species we identified. We hope that by providing a more accurate set of read counts for the same TCGA samples, our dataset can serve as a valuable scientific resource, enabling researchers to better distinguish genuine microbial signals from background noise and contaminants in future investigations.

We also extended our previous work to add comparisons to the fungal genomes found in cancer by Narunsky-Haziza et al. *(4)*. Upon looking at the data published in that study, we discovered that several of the key fungal species used to create cancer-specific signatures had excessively high read counts, and that at least some of those counts were the result of contamination in the fungal genome sequences. The contaminants included both human DNA and vector contaminants, either of which can lead to high numbers of false positives when doing metagenomic analysis.

Multiple other studies have recently reported microbiomes in cancer, including a widely publicized report of a fungal mycobiome in pancreatic cancer *(30)* that was only recently (in 2023) shown to be flawed *(31)*, likely due to mis-identification of fungal reads in the original data. Similarly, a 2020 study reported tumor type-specific microbes in seven types of cancer *(32)*, but an attempt to replicate those findings in breast cancer found no evidence of microbes at all *(33)*. Taken together with the results described here, these reports suggest that claims regarding microbiomes and cancer need to be scrutinized more rigorously than they have been in the recent past, and that contamination of human samples with environmental microbes can easily be mistaken for a genuine signal.

Finally, we did serendipitously find a number of individual microbes present in high abundance in some samples. Some of these were consistent with prior knowledge, while others may be worthy of further investigation, ideally with additional, independent experimental data.

## Methods

We downloaded sequence data from the Genome Data Commons at the U.S. National Cancer Institute (gdc.cancer.gov) for 25 cancer types from the TCGA project. Metadata on all samples used in this study and previous studies, including unique sample identifiers from TCGA, can be found in **Table S13**. In total, data from 5,734 WGS samples were downloaded from the NCI data portal, which had aligned the reads to either hg19 or GRCh38 using bwa *(8)*. Of these samples, 2824 represented solid tumors, 569 were solid normal tissue, 64 were blood-derived cancer, and 2277 were blood-derived normal. We downloaded all reads that were unmapped by TCGA to their reference genomes, which included the human genome (either hg19 or GRCh38, depending on the date of data collection) as well as human papilloma virus (multiple subtypes), hepatitis C virus 1 and 2, hepatitis B, and Epstein-Barr virus. We then aligned the unmapped reads against the CHM13 human reference using Bowtie2 *(16)* to remove more human reads (**Table 1**).

We classified these two-pass filtered read sets using KrakenUniq *(7)* using its default parameters in paired-read mode, which treats each pair of reads as a single discontiguous sequence. We classified the ∼15 billion two-pass filtered read pairs (Table 1) against a curated database, Microbial2023, containing all RefSeq bacterial, archaeal, and viral complete genomes, a collection of curated eukaryotic pathogen genomes from EuPathDB54, 10,798 vector sequences from UniVec and EmVec, the CHM13v2.0 genome, and the GRCh38.p14 human genome. In total, Microbial2023 contains 50,079 genomes (29,798 species) divided into 34,452 bacteria (16,150 species), 14,018 viruses (12,910 species), 534 archaea (422 species) and 503 eukaryotic microbes (316 species). The list of species and their GenBank accessions can be found in **Table S14**. The Microbial2023 database, which is 535GB in size, can be downloaded from https://benlangmead.github.io/aws-indexes/k2. A total of 891,242,697 non-vector, non-human reads were classified by KrakenUniq at the genus level or below; of these, 827,851,151 were classified at the species level or below. Note that ∼63.4 million reads were classified at the genus level but not at the species level because they matched more than one species equally well.

In order to conduct a more comprehensive search against fungi, we created a separate database containing all 572 fungal genomes (557 species) in the NCBI RefSeq database as of mid-2023, which we designate as Fungi_RefSeq. The list of species and their GenBank accessions can be found in **Table S15**. After processing each sample using Microbial2023, we extracted all reads that were either failed to match or were classified as fungi (taxid=4751), and screened them against Fungi_RefSeq using KrakenUniq with default parameters. These steps yield read counts for all 5,734 samples against the 557 fungal species.

To compare our bacterial, archaeal, and viral read counts to Poore *et al*.’s results, we compared the sample metadata and species and determined that 4,550 out of 5,734 TCGA samples were analyzed in both studies, and 1,289 out of 1,993 microbial genera were reported in both studies. (Note that because Microbial2023 is a newer database, it has some species missing from the Poore *et al*. study. Conversely, because Microbial2023 only includes finished genomes while Poore *et al*. included draft genomes, some species and genera used in Poore *et al*. are missing from Microbial2023.) Similarly for Narunsky-Haziza *et al*.’s results, we matched 4,271 out of 5,734 TCGA samples. All 224 fungal species used in Narunsky-Haziza *et al*. were included in our re-analysis. A list of species name changes from RefSeq200 to RefSeq220 involving some of these fungi can be found in **Table S16**.

In our analysis of the high read counts in *M. globosa, T. asahii, R. collo-cygni*, and *C. albicans*, we aligned human-filtered reads from the samples against their respective reference genomes using the same Bowtie2 parameters as used by Narunsky-Haziza *et al*. (--end-to-end --very-sensitive -k 16 --np 1 --mp 1,1 --rdg 0,1 --rfg 0,1 --score-min L,0,-0.05).

## Acknowledgements

The authors wish to thank David Lipman, Mihaela Pertea, Ales Varabyou, and Aleksey Zimin for helpful comments on earlier drafts of this manuscript.

## Author contributions

Conceptualization: YG, SLS

Methodology: YG, JL, MR, DP, SLS

Data Analysis: YG, JL, MR, DP, SLS

Funding acquisition: SLS

Supervision: SLS

Writing – original draft: YG, JL, MR, DP, SLS

Writing – review & editing: YG, SLS

## Data Availability

All supplemental tables and files from this study are freely available at https://github.com/yge15/TCGA_Microbial_Content.

