## Supplementary material for "Comprehensive analysis of microbial content in whole-genome sequencing samples from The Cancer Genome Atlas project": TableS8

**Supplementary Table SX. TCGA Study Abbreviations**

| **Study Abbreviation** | **Study Name** |
| --- | --- |
| LAML | Acute Myeloid Leukemia |
| ACC | Adrenocortical carcinoma |
| BLCA | Bladder Urothelial Carcinoma |
| LGG | Brain Lower Grade Glioma |
| BRCA | Breast invasive carcinoma |
| CESC | Cervical squamous cell carcinoma and endocervical adenocarcinoma |
| CHOL | Cholangiocarcinoma |
| LCML | Chronic Myelogenous Leukemia |
| COAD | Colon adenocarcinoma |
| CNTL | Controls |
| ESCA | Esophageal carcinoma |
| FPPP | FFPE Pilot Phase II |
| GBM | Glioblastoma multiforme |
| HNSC | Head and Neck squamous cell carcinoma |
| KICH | Kidney Chromophobe |
| KIRC | Kidney renal clear cell carcinoma |
| KIRP | Kidney renal papillary cell carcinoma |
| LIHC | Liver hepatocellular carcinoma |
| LUAD | Lung adenocarcinoma |
| LUSC | Lung squamous cell carcinoma |
| DLBC | Lymphoid Neoplasm Diffuse Large B-cell Lymphoma |
| MESO | Mesothelioma |
| MISC | Miscellaneous |
| OV | Ovarian serous cystadenocarcinoma |
| PAAD | Pancreatic adenocarcinoma |
| PCPG | Pheochromocytoma and Paraganglioma |
| PRAD | Prostate adenocarcinoma |
| READ | Rectum adenocarcinoma |
| SARC | Sarcoma |
| SKCM | Skin Cutaneous Melanoma |
| STAD | Stomach adenocarcinoma |
| TGCT | Testicular Germ Cell Tumors |
| THYM | Thymoma |
| THCA | Thyroid carcinoma |
| UCS | Uterine Carcinosarcoma |
| UCEC | Uterine Corpus Endometrial Carcinoma |
| UVM | Uveal Melanoma |
