## Supplementary material for "Comprehensive analysis of microbial content in whole-genome sequencing samples from The Cancer Genome Atlas project": FigureS1

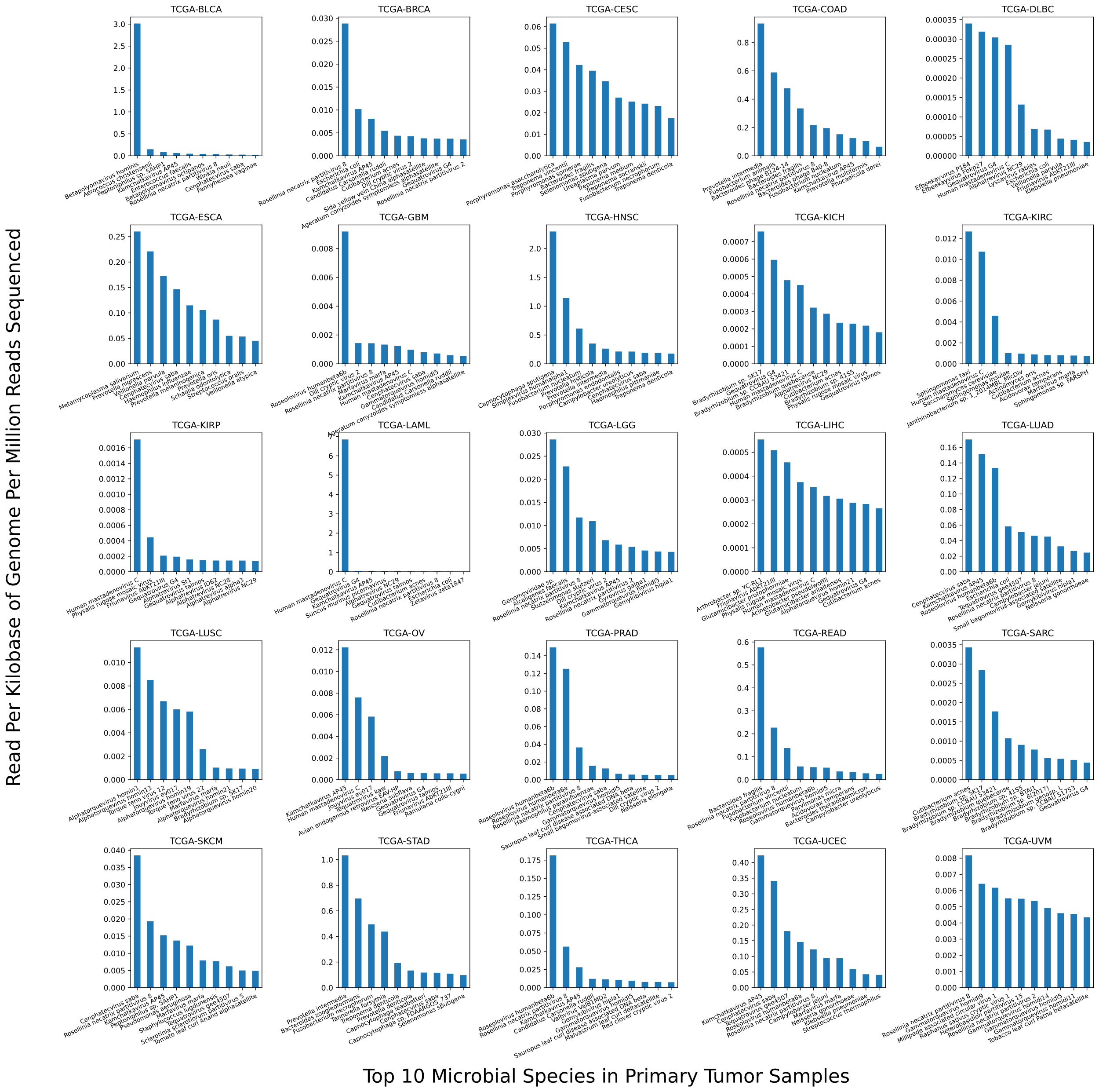

**Supplementary Figure S1.** Normalized counts of the top 10 microbial species in primary tumor samples for each of 25 cancer types. The X-axis shows the species names, sorted by maximum normalized counts, measured as reads per kilobase of genome per million reads sequenced.
